# MR1-restricted T cell clonotypes are associated with ‘resistance’ to *M.tuberculosis* infection

**DOI:** 10.1101/2022.10.12.511825

**Authors:** Deborah L. Cross, Erik D. Layton, Krystle K.Q. Yu, Malisa T. Smith, Melissa S. Aguilar, Shamin Li, Harriet Mayanja-Kizza, Catherine M. Stein, W. Henry Boom, Thomas R. Hawn, Philip Bradley, Evan Newell, Chetan Seshadri

## Abstract

T cells are required for a protective immune response against the human adapted pathogen *Mycobacterium tuberculosis* (M.tb). We recently described a cohort of Ugandan household contacts of tuberculosis cases that appear to ‘resist’ M.tb infection (RSTRs) and showed that these individuals harbor IFN-γ independent T cell responses to M.tb-specific peptide antigens. However, T cells also recognize non-protein antigens via antigen presenting systems that are independent of genetic background, leading to their designation as donor-unrestricted T (DURT) cells. We used combinatorial tetramer staining and multi-parameter flow cytometry to comprehensively characterize the association between DURTs and ‘resistance’ to M.tb infection. We did not observe a difference in peripheral blood frequencies of invariant natural killer T (iNKT) cells, germline encoded mycolyl-reactive (GEM) T cells, or γδ T cells between RSTRs and matched controls with latent M.tb infection (LTBIs). However, we did observe a 1.65-fold increase in frequency of circulating MR1-restricted T (MR1T) cells among RSTRs in comparison with LTBI (p=0.03). Multi-modal single cell RNA-sequencing of 18,251 MR1T cells sorted from a subset of donors revealed 5150 clonotypes that expressed a common transcriptional program, the majority of which were private. Deep sequencing of the TCR-α repertoire revealed several DURT clonotypes that were expanded among RSTRs, including at least two MR1T clonotypes. Taken together, our data reveal unexpected donor-specific diversity in the TCR repertoire of human MR1T cells as well as associations between MR1 clonotypes and ‘resistance’ to M.tb infection.

## INTRODUCTION

*Mycobacterium tuberculosis* (*M.tb*) is the aetiological cause of tuberculosis (TB) which caused an estimated 1.5 million deaths worldwide in 2020^1^. Nearly 1.7 billion people who have been exposed to *M.tb* are clinically asymptomatic but may harbor a ‘latent’ TB infection (LTBI) on the basis of a positive tuberculin skin test (TST) or IFN-γ release assay (IGRA)^2,3^. Most animal models seek to recapitulate active disease rather than latent M.tb infection, so the immune mechanisms underlying protection from *M.tb* infection rather than disease are currently undefined^4^. We recently described a cohort of Ugandan household contacts of active TB cases who fail to convert their TST or IGRA despite high risk of exposure to *M.tb* ^5,6^. We further showed that these individuals have primed IFN-γ independent T cell responses to *M.tb*-specific antigens, confirming the epidemiologic evidence of exposure^7^. Since these individuals also did not develop active TB over a median of 9.5 years of follow up, we hypothesize that they ‘resist’ *M.tb* infection (RSTRs). Whether other T cell profiles are also associated with the RSTR phenotype is currently unknown.

T cells typically recognize foreign peptide antigens through a genetically rearranged T cell receptor (TCR) when bound to highly polymorphic major histocompatibility complex (MHC) molecules^8^. T cells are also capable of recognizing non-peptide antigens through MHC-independent antigen presentation systems^9^. For example, T cells are activated by lipids and small molecules presented by cluster of differentiation 1 (CD1) and major histocompatibility complex (MHC)-related protein 1 (MR1), respectively^10,11^. Further, T cells with a γδ T cell receptor (TCR) can recognize non-peptide antigens presented by butyrophilin molecules, CD1, and MR1^12–14^. Because CD1, MR1, and butyrophilin exhibit limited sequence diversity, the T cells that act through these systems are called donor-unrestricted T cells (DURTs)^15^.

MR1-restricted T-cells (MR1T-cells) are strongly activated by derivatives of the riboflavin biosynthesis pathway and capable of lysing bacterially infected cells^11,16^. Mucosal associated invariant T (MAIT) cells are a subset of MR1Ts that are abundant in human peripheral blood and characterized by expression of a germline-encoded TCR-α chain utilizing TRAV1-2 and either TRAJ33, TRAJ20 or TRAJ12 gene segments^10,17,18^. X-ray crystallography studies have demonstrated the importance of the TCR-α chain in recognition of MR1-presented antigens with additional reports demonstrating a role for TRBV genes in differential bacterial sensing^11,19–22^. In addition, there is a growing appreciation of MR1T-cells that lack the conserved TCR gene usage that characterizes MAIT cells and may mediate recognition of alternative ligands^23^. For example, photolumazines were recently identified as a novel class of mycobacterial-derived ligands that could be presented by MR1 to MAIT cells^24^.

Here, we sought to comprehensively study the association between DURTs and ‘resistance’ to *M.tb* infection. We found that the frequency of circulating MR1T cells was increased in RSTRs when compared to matched LTBI controls. This result led to a detailed study of MR1T cells using multi-modal single-cell RNA-sequencing. We found that over 80% of MR1T clonotypes could only be detected in a single donor, highlighting an unexpected donor specificity of this DURT subset. At least two of these MR1T clonotypes, but not the canonical MAIT TCR-α, were preferentially detected in the blood of RSTRs. A broader survey of TCR-α clonotypes defined by immunosequencing also revealed preferential clone sharing among RSTRs when compared to LTBI controls. Together, these data reveal associations between MR1-restricted as well as other DURT clonotypes of unknown specificity with ‘resistance’ to *M.tb* infection.

## METHODS

### Study Participants

The full details of the parent clinical study, including enrollment and sample collection, have been previously described^6^. Briefly, household contacts of sputum culture-positive cases of pulmonary TB were enrolled as part of the Kawempe Community Health Study conducted between 2002 and 2012. Contacts were sputum culture-negative at enrollment and had no radiological evidence of active *M.tb* infection. Enrolled individuals were longitudinally profiled for signs of latent M.tb infection by tuberculin skin test (TST) (Mantoux method, 0.1□ml of 5□tuberculin units of PPD, Tubersol; Connaught Laboratories) over a 2-year observation period. TST positivity was defined as an induration of > 10 mm for HIV-individuals and > 5mm for HIV+ individuals. Overall, 2,585 individuals were enrolled and 10.7% of this group (n□=□198) were persistently tuberculin skin test (TST) negative over a two-year follow-up. Between 2014 and 2017, 691 individuals from this original household contact study were retraced, of which 441 (63.8%) were successfully reidentified and willing to participate in a subsequent longitudinal follow-up study. The median time between the initial study and retracing was 9.5 years. Retraced individuals completed three QuantiFERON-TB Gold (QFT) assays over two years. At their final visit, individuals were also tested by TST. A definite classification was assigned if TST assay results (five from the original study and one at the end of the re-tracing study) and three QFTs from the retracing study were concordantly negative or positive (RSTR or LTBI, respectively). Peripheral blood mononuclear cells (PBMCs) were isolated from whole blood by Ficoll-Hypaque density centrifugation and cryopreserved until use.

### Ethics

The retracing study protocol was approved by the National AIDS Research Committee, The Uganda National Council on Science and Technology, and the institutional review board at University Hospitals Cleveland Medical Center. All study participants gave written, informed consent, approved by the institutional review boards of the participating institutions.

### Generation of Tetramers

All tetramers were generated as published previously^25,26^. Biotinylated CD1b monomers were provided by the National Institutes of Health Tetramer Core Facility (Emory University, Atlanta, GA). Glucose monomycolate (GMM) was a generous gift from Dr. Branch Moody and synthetic diacylated sulfoglycolipid (Ac_2_SGL was provided by Professor Adriaan Minnaard. For CD1b monomer loading, GMM or Ac_2_SGL were dried down under a nitrogen stream and then sonicated into 50 mM sodium citrate buffer at pH 4, containing 0.25% with 3-[(3-cholamidopropyl) dimethylammonio]-1-propanesulfonate (CHAPS) (Sigma, St. Louis, MO) for two minutes at 37°C. CD1b monomer was added with the resulting lipid suspensions at either 100-fold or 40-fold molar excess of CD1b monomer. The monomer and lipid suspensions were subsequently incubated at 37°C for 2 hours, vortexing every 30 minutes. Following incubation, the solution was neutralized to pH 7.4 using 6 uL of 1 M Tris pH 9. Loaded lipid monomers were tetramerized by the addition of 1.25 molar equivalents of fluorophore-conjugated streptavidin (either BV421 conjugated for CD1b-GMM (BioLegend, San Diego, CA) or PE-conjugated for Ac2GSL (Life Technologies, Carlsbad, CA)), assuming a 4:1 ratio of biotin to streptavidin was needed. Streptavidin was added over the course of 2 hours, 1/10th of the needed volume streptavidin was added, the vial was mixed, and then incubated for 10 minutes before another volume was added. Tetramers were filtered for aggregates through a SpinX column (Sigma, St. Louis, MO) then stored at 4°C until use. Mock-loaded CD1b tetramers were generated by an analogous process without the addition of exogenous lipids.

PBS-57 (α-galactosylceramide or α-GalCer)-loaded and mock-loaded human CD1d monomers were provided by the National Institutes of Health Tetramer Core Facility (Emory University, Atlanta, GA). Tetramers were prepared as previously described^27^. Briefly, 10 uL of stock 2 mg/mL CD1d-α-GalCer was combined with 2.6 uL of streptavidin BV650 every 10 minutes for 100 minutes until a final volume of 26 uL was reached. The tetramer was filtered through a SpinX column (Sigma, St. Louis, MO) to remove aggregates and then stored at 4°C until use. MR1-5-(2-oxopropylideneamino)-6-d-ribitylaminouracil (5-OP-RU) tetramer was obtained from the National Institutes of Health Tetramer Core Facility (Emory University, Atlanta, GA) and used as provided.

### Flow cytometry

PBMC were thawed and washed in warmed RPMI 1640 (Gibco, Waltham, MA) supplemented with 10% fetal calf serum (FBS) (Hyclone, Logan, UT) and 2 uL/mL Benzonase (Millipore, Burlington, MA). PBMC were then resuspended at a density of two million cells/mL in RPMI/10% FBS and allowed to rest overnight at 37°C in humidified incubators supplemented with 5% CO_2_. The following day, the PBMC were enumerated using the Guava easyCyte. One million cells/well were plated into a 96-well U-bottom plate with up to four wells plated per sample. Cells were blocked with human serum (Valley Biomedical, Winchester, VA) prepared in FACS buffer (1x PBS (Gibco, Waltham, MA) supplemented with 0.2% BSA (Sigma, St. Louis, MO)) mixed 1:1 for 15 minutes at 4°C. A 13-color multiparameter flow cytometry panel targeting donor-unrestricted T cell populations, lineage, and memory populations was used to characterize cells from each individual. Cells were centrifuged at 1800 rpm for 3 minutes, resuspended in 50 uL of FACS buffer containing CD1b-GMM, CD1b-AM Ac_2_SGL, CD1d-□-GalCer, MR1-5-OP-RU, and mock-loaded tetramers and incubated at room temperature for 60 minutes. The cells were then washed twice with PBS and stained with Live/Dead Fixable Green Dead Cell Stain Kit (Life Technologies, Carlsbad, CA) per the manufacturer’s instructions. Following a 15 minute incubation at room temperature, the cells were washed twice in PBS and then labelled with anti-CD3 ECD (clone UCHT1; Beckman Coulter, Brea, CA), anti-CD4 APC Alx750 (clone 13B8.2; Beckman Coulter, Brea, CA), anti-CD8α PerCP Cy5.5 (clone SK1; BD Biosciences, San Jose, CA), anti-CD45RA BUV737 (clone HI100; BD Biosciences, San Jose, CA), anti-CCR7 BUV395 (clone 150503; BD Biosciences, San Jose, CA), anti-TRAV1-2 BV510 (clone 3C10; Biolegend, San Diego, CA), anti-Pan-gδ PE-Vio770 (clone 11f2; Miltenyi Biotech, Bergisch Gladbach, Germany) and anti-Vδ2 BV711 (clone B6; Biolegend, San Diego, CA) for 60 minutes at room temperature. The optimal titers of all antibodies and tetramers were determined prior to use. After two final washes in FACS buffer, the cells were fixed in 1% paraformaldehyde (Electron Microscopy Sciences, Hatfield, PA) and acquired on a BD LSRFortessa (BD Biosciences, San Jose, CA) equipped with blue (488 nm), green (532 nm), red (628 nm), violet (405 nm), and ultraviolet (355 nm) lasers using standardized good clinical laboratory practice (GCLP) procedures to minimize the variability of data generated.

### Cell Sorting

PBMCs were thawed in warmed RPMI (10% FBS with 2uL/mL benzonase) and washed twice in 5mL at 1200rpm for 10 minutes. After washing, cells were resuspended in 96-well U-bottom plates and blocked prior to staining with 1 μg/mL anti–CD40 (clone HB14, Miltenyi Biotec) for 30 minutes at 37°C. Following blocking, cells were stained with MR1-5-OP-RU tetramer as described above for 60 minutes at room temperature. Samples were washed in FACS buffer to remove unbound tetramer prior to surface staining with anti-TCR□β APC (clone IP26) and anti–CD7 FITC (clone CD7-6B7, Biolegend, San Diego, CA) for 30 minutes at 4°C. Samples were washed and resuspended in 500uL of FACs buffer prior to sorting using the BD FACSAria II Cell Sorter equipped with blue (488 nm), red (641 nm), and violet (407 nm) lasers. CD7+/TCRβ+/MR1-5-OP-RU+ live intact singlets were sorted for downstream analysis (**Supplementary Figure 1A**).

### CITE-Seq and scTCR-seq

For CITE-Seq experiments, cells were stained with oligo-tagged antibodies: anti-CD3 (clone UCHT1), anti-CD4 (clone RPA-T4), anti-CD8a (clone RPA-T8), anti-CD56 (clone QA17A16), anti-CD45RO (clone UCHL1), anti-CD69 (clone FN50), anti-CD27 (clone O323), anti-CD39 (clone A1), anti-CD244 (clone C1.7), anti-CD127 (IL-7Ra; clone A019D5), anti-CD38 (clone HIT2), anti-CD71 (clone CY1G4), anti-CXCR3 (clone G025H7), anti-CD196 (clone G034E3), anti-CD161 (clone HP-3G10), anti-HLA-DR (clone L243), anti-CX3CR1 (clone K0124E1), anti-CD81 (clone 5A6), anti-CD28 (clone CD28.2), anti-CD26 (clone BA5b), anti-TCR Va7.2 (clone C0581), anti-Human Hashtag 1 (CD45; clone LNH-94), anti-Human Hashtag 2 (CD45; clone LNH-94), anti-Hashtag 3 (CD45; clone LNH-94) and anti-Hashtag 4 (CD45; clone LNH-94). All TotalSeq antibodies were obtained from Biolegend, San Diego, CA. Cells were stained with the sorting antibody cocktail and MR1-5-OP-RU tetramer for 30 minutes at 4°C and sorted as described above.

Single cell library preparation for surface protein, mRNA and TCRs was performed using the Chromium Single Cell V(D)J Reagent Kit v1.1 (10x Genomics, Pleasanton, CA). The TCR V(D)J region was specifically enriched for a separate library preparation using the Chromium Single-Cell V(D)J Enrichment Kit (10x Genomics, Pleasanton, CA). Cell suspensions were combined with barcoded, single-cell 5’ gel beads and loaded onto Chromium Next GEM Chip G (10x Genomics, Pleasanton, CA) at a limiting dilution such that a single bead and a single cell are partitioned into a sphere. Libraries were sequenced by Illumina sequencing (NextSeq 500/550 platform). Sequence alignment was performed using Cell Ranger (10x Genomics, Pleasanton, CA). All mRNA and V(D)J reads were aligned to the GRCh38 human reference genome.

### TCR-α/δ (TCRAD) Immunosequencing

For each sample (n = 40), DNA was extracted from PBMCs using the Qiagen DNeasy Blood and Tissue kit (Qiagen, Hilden, Germany). DNA was quantified using TapeStation (Agilent, Santa Clara, CA). TCR-α/δ chains were sequenced using the ImmunoSEQ high-throughput sequencing platform (Adaptive Biotechnologies, Seattle, WA)^28^. For each sample, multiplex PCR was used to amplify rearranged VDJ sequences followed by high throughput sequencing using Illumina technologies (Illumina, San Diego, CA). PCR amplification bias was minimized by internal controls in the ImmunoSEQ assay^29^. Raw data were exported to R from the Adaptive ImmunoSEQ Analyser.

### Data Analysis

#### Flow Cytometry

Spillover compensation and initial quality assessment were performed using FlowJo version 9.9.6 (FlowJo, TreeStar Inc.). **Supplementary Figure 1B** provides a representative gating strategy for identification of donor-unrestricted T-cell subsets. T-cell subset counts, as a proportion of either CD3+ live, intact singlets or CD3+/MR1-5-OP-RU+ live, intact singlets, were exported and analyzed in the R programming environment. All samples had acceptable viability and frequencies of CD3+ events and were included in downstream analysis. Subset frequencies were compared between groups using a Wilcoxon Rank Sum test.

#### CITE-Seq

All data analysis was performed in the R programming environment using *Seurat* workflows. Hashtag oligos (HTO) were transformed using a centered log-ratio transformation (CLR) applied with the *ScaleData* function in *Seurat 4.0*^30^. Cells were demultiplexed into original donor samples based on enrichment of HTOs using the *HTODemux* function in *Seurat*. Next, the probability of a cell being degraded was estimated using a probabilistic mixture model framework implemented using the *miQC* package^31^. Percentage of mitochondrial reads and total gene count per cell were used as input variables. A posterior probability threshold of 0.75 was used to remove compromised cells. Initial analyses of each sequencing batch identified batch effects in both gene and protein expression datasets. Batches were integrated using canonical correlation analysis (CCA) in *Seurat* prior to further downstream analysis^32^. Variability in gene expression sequencing depth between samples was corrected for by normalization using the function *scTransform* in *Seurat*^33^. ADT counts for each marker were transformed using CLR prior to downstream analysis. Gene and surface protein expression were used to define clusters using the weighted nearest neighbors workflow in *Seurat* (WNN, resolution = 0.5)^30^. Differential expression of genes between clusters was performed in *Seurat* using Wilcoxon Rank Sum Tests. These tests were performed on the RNA assay using *FindConservedMarkers* in Seurat using batch as a grouping variable. A gene was differentially expressed for a given cluster if it was significantly changed between that cluster and the remaining data (FDR = 0.05 with Bonferroni correction for multiple testing). Highly variable genes in the dataset (default selection of top 3000 genes was used) were used as input for principal component analysis (PCA). The default number of principal components was computed (n = 50). Other dimensionality reductions such as UMAP were performed using principal components as input. UMAP and PCA were both implemented in *Seurat*.

#### scTCR-seq

Reads were mapped to the *TRA* and *TRB* loci using CellRanger software. Contigs were filtered for productive rearrangements with in-frame CDR3 sequences. Additional filtering was applied to include only viable cells (as identified in scRNA-seq pre-processing) and barcodes with a paired TCR-α and TCR-β chain. Finally, contigs were collapsed to produce a data frame where each row represented a single barcode using the *scRepertoire* package^34^. Global repertoire metrics (evenness, diversity, number of unique clonotypes) were computed in R using the *Immunarch* package^35^. Identification of biochemically similar TCRs was performed using *TCRdist* in Python^36,37^. In TCRdist, CDR loops that contact the pMHC complex are concatenated into a single string. These are then used as input to compute a distance matrix between all unique TCR clonotypes in our dataset. The distance measure is a similarity-weighted Hamming distance with a gap penalty applied in cases where CDR sequences are different lengths. The CDR3 sequence is given a higher weight given its importance in pMHC recognition. The TCRdist distance matrix was then used for clustering (hierarchical clustering using the *hclust* function in the *stats* package v3.6.1 in base R) and dimensionality reduction (multidimensional scaling using the *cmdscale* function in the *stats* package).

#### Graph-vs-graph analysis

Relationships between single-cell gene expression profile and TCR clonotype were assessed using clonotype neighbor graph analysis (CoNGA)^38^. Briefly, similarity graphs were constructed in gene expression and TCR space respectively. In the gene expression graph, nodes represented cells and edges represented correlation in gene expression. In the TCR graph, nodes represented cells and edges represented pairwise TCR similarity based on the TCRdist distance metric (described above). Only cells that passed pre-processing described above were included in the analysis. Neighborhood overlap between the two graphs was evaluated. The number of vertices connected to each clonotype in the gene expression and TCR graphs were individually counted and a significance score was generated based on whether the number of overlapping connections was greater than expected by chance.

#### TCR-α/δ Immunosequencing

Raw data was exported from Adaptive ImmunoSEQ Analyser and analysed in R. Clonotypes that were in-frame, had both a V-gene and a J-gene assignment and with a CDR3α amino acid length of more than 5 were included in further analyses. Following pre-processing, contingency tables of TCR-α or TCR-δ clonotypes were statistically assessed for enrichment in either RSTR or LTBI donors using two-sided Fisher’s exact tests and a nominal p-value cutoff of 0.01. In addition, significantly enriched clonotypes identified in immunosequencing data were parsed for known MR1T clonotypes using our scTCR-sequencing as a reference. Clonotype matching based on nucleotide sequence yielded no hits so matches based on CDR3α amino acid sequence and TCR-α V- and TCR-α J-gene usage were used instead.

## RESULTS

### MR1-restricted T-cells are expanded in the peripheral blood of RSTRs compared to LTBI controls

We leveraged samples collected as part of a longitudinal study of TB household contacts in Uganda, as we have previously reported^5,6^. Three sequential IGRAs, measured by QuantiFERON-TB Gold, were performed on blood samples and one additional TST was performed as part of this study that took place approximately 9.5 years after the initial *M.tb* exposure^6^. Human immunodeficiency virus (HIV)-negative subjects who remained concordantly negative for all tests were defined as ‘resisters’ (RSTRs) and control subjects with LTBI were defined by consistently positive results at all time points by both IGRA and TST. For this analysis, we selected a representative subset of RSTRs and LTBI controls after matching by age (≥15 years), gender, and epidemiologic risk score (**Supplementary Table 1**). We used combinatorial tetramer staining and multi-parameter flow cytometry to quantify the frequencies of donor-unrestricted T-cells in RSTRs (n=25) and LTBIs (n=25)^26^ (**Figure 1A, Supplementary Figure 1B**). We observed no difference in circulating frequencies of γδ T cells, iNKT cells, GMM-specific, or Ac_2_SGL-specific T cells between RSTRs and LTBI controls (**Figure 1B**). However, circulating frequencies of CD3+ MR1-5-OP-RU+ T-cells (MR1T-cells) were 1.65-fold higher in RSTR when compared to LTBI controls (**Figure 1C**, p = 0.028). Frequencies of subsets defined by TRAV1-2 or co-receptor (CD4 or CD8) expression all showed a similar trend toward expansion in RSTRs, suggesting that no single subset was responsible for the difference in total MR1Ts **(Figure 1D)**. Taken together, these data highlight the association between circulating frequencies of MR1Ts and ‘resistance’ to *M.tb* infection.

**Figure 1.**
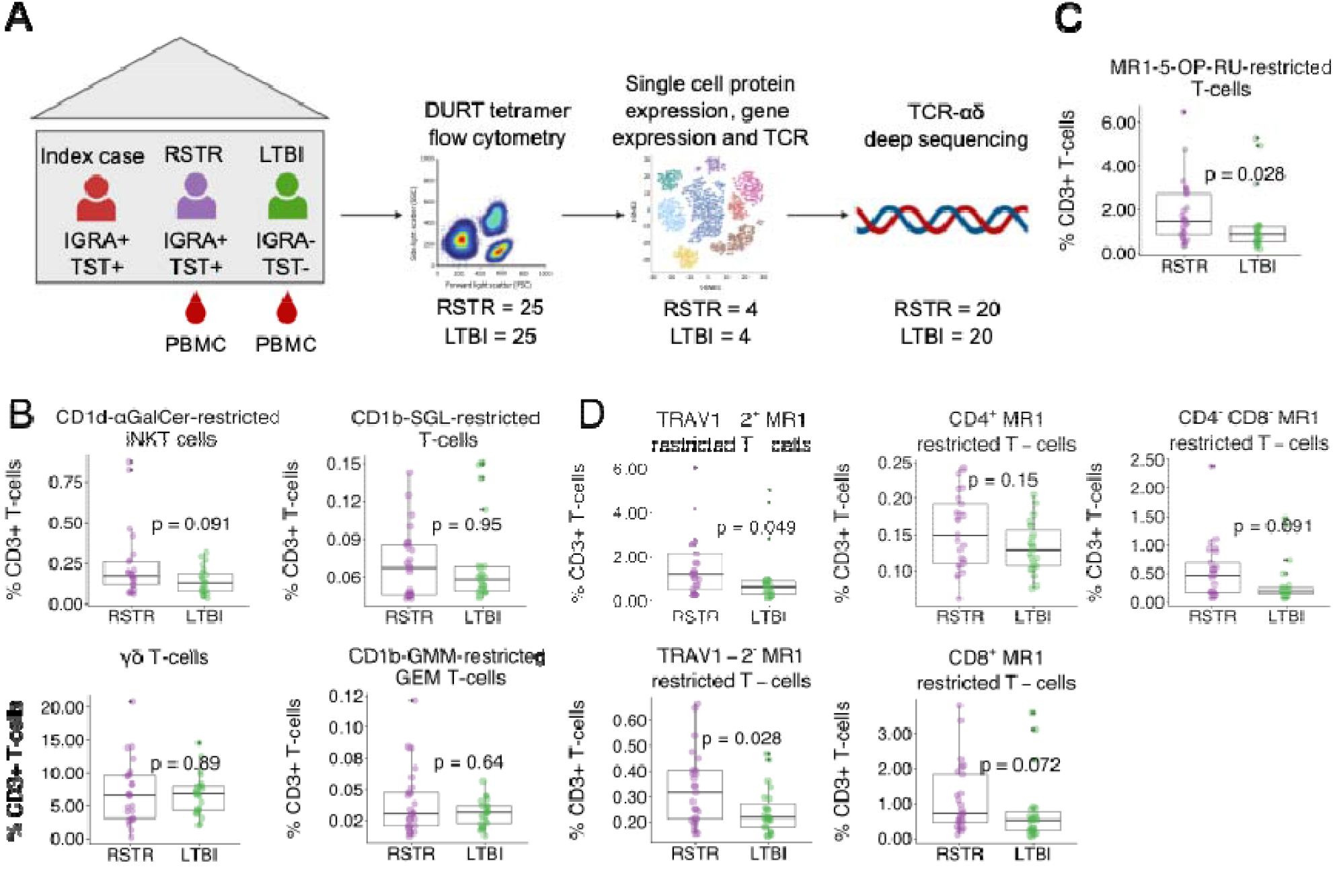
MR1-restricted T-cells are expanded in the peripheral blood of RSTRs compared to LTBI controls. (A) Overview of study design. Household contacts of active TB cases were classified as RSTR or LTBI as defined by longitudinal testing with TST and IGRA. Combinatorial tetramer staining and multi-parameter flow cytometry was used to quantify frequencies of donor-unrestricted T cells in RSTRs (n=25) and LTBIs (n=25). MR1-5-OP-RU tetramer was used to sort MR1T cells for multi-modal single-cell RNA-sequencing from RSTRs (n=4) and LTBIs (n=4), including identification of TCR clonotype, transcriptional profiling, and surface protein expression with CITE-Seq. Deep-sequencing of the TCR-α and TCR-δ repertoire was performed using the ImmunoSEQ platform (Adaptive Biotechnologies) from RSTRs (n=20) and LTBIs (n=20). (B) Frequencies of CD1-restricted and γδ T cells stratified by group as a proportion of live CD3+ T-cells. (C) Frequencies of MR1-5-OP-RU staining T-cells as a proportion of live CD3+ T-cells stratified by group (D) Among MR1–OP-RU staining T cells, the frequencies of T cells expressing CD4, CD8, or TRAV1-2 as a proportion of live CD3+ T-cells stratified by group. Statistical testing was performed using the Wilcoxon rank sums test and unadjusted p-values are displayed.

### The majority of the MR1-restricted T cell receptor repertoire consists of donor-specific clonotypes

Clonotypic diversity in the MR1T compartment is generally thought to be low given the majority of clonotypes express a semi-invariant T cell receptor (TCR)^23^. However, evidence of MR1 T-cell-restricted populations with greater TCR diversity has been reported^39^. Therefore, to determine whether MR1T clonotypic diversity is associated with *M.tb* resistance, we sorted MR1-5-OP-RU+ T-cells from RSTRs (n=4) and LTBI (n=4) and performed multi-modal scRNA-Seq on 70,535 MR1T-cells. Following pre-processing, 18,251 cells with paired scTCR-seq information were included in downstream analysis (**Supplementary Figure 2A**). Across all donors (n = 8), 5150 TCR clonotypes were identified, defined by both CDR3-α and CDR3-β nucleotide sequences. The number of unique clonotypes, overall richness, and repertoire clonality were similar between RSTRs and LTBIs (**Supplementary Figure 2B**). Consistent with the prior literature, we found that the majority of the MR1T repertoire was composed of expanded clonotypes that appeared at least twice in our dataset^40^. On average, expanded clonotypes represented 75% of the MR1T repertoire (**Figure 2A**). Surprisingly, we also observed a significant minor fraction (∼25%) of each donor’s repertoire that was unexpanded, appearing only once in the repertoire. The proportion of unexpanded (n = 1), expanded (1 < n <= 50) and hyperexpanded (n > 50) MR1 T cell clonotypes were highly variable between donors (**Supplementary Figure 2C**) but few differences in proportions were noted between RSTR and LTBI (**Figure 2B**).

**Figure 2.**
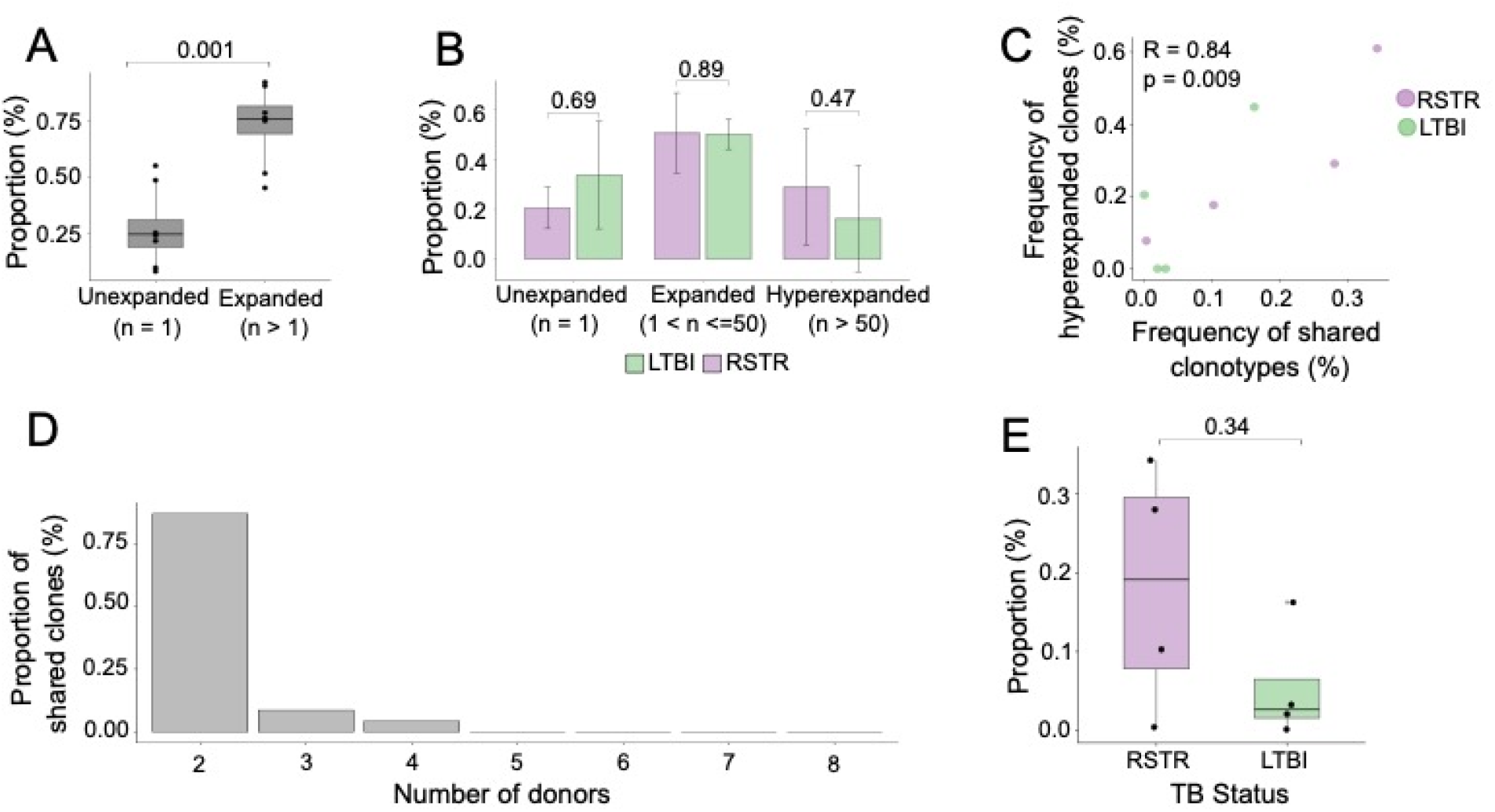
The majority of the MR1-restricted T cell receptor repertoire consists of donor-specific clonotypes. (A) The proportion of unexpanded (n = 1) and expanded (n > 1) MR1T clonotypes as defined by nucleotide sequence is displayed as a percentage of each donor’s total repertoire (n=8). (B) The proportion of unexpanded (n = 1), expanded (1 < n <= 50) and hyperexpanded (n > 50) MR1T clonotypes as a percentage of the total repertoire is displayed stratified by group. Height of the bar reflects the mean and error bars represent standard deviation. (C) Pearson correlation between the frequency of hyperexpanded clonotypes and frequency of public clonotypes. Each point represents one donor and points are coloured according to group. (D) Among shared clones, the proportion that are detected in two or more donors is shown. (E) The proportion of shared MR1T clonotypes is displayed stratified by group. Each point is an individual donor. Statistical testing was performed using the Wilcoxon rank sums test and unadjusted p-values are displayed.

Given the extent of clonal expansion and the designation of MR1Ts as DURTs, we hypothesized that the majority of expanded clonotypes might be shared between donors. Frequencies of shared clonotypes were strongly correlated with the frequency of hyperexpanded clonotypes (R = 0.84, p = 0.010) suggesting shared clonotypes represented a major fraction of the total repertoire (**Figure 2C**). To our surprise, only 23 of 5150 unique clonotypes in our dataset were shared among two or more donors, with 20 (87%) of 23 clonotypes being shared only between two individuals (**Figure 2D**). We found no clonotypes that were common to all donors studied. Finally, we found that the frequency of shared clonotypes trended higher in RSTRs compared to LTBI donors (p = 0.34) (**Figure 2E**). Taken together, these data reveal higher than expected donor restriction in the MR1T repertoire with only a minor proportion of highly expanded clonotypes being shared between donors.

### MR1T clonotype diversity is constrained by TCR gene usage and CDR3 length

We next sought to gain further insight into the features that distinguish shared from donor-specific MR1T clonotypes. In both compartments, the majority of clonotypes expressed TRAV1-2, consistent with published data and validating our sorting strategy^20^ (**Figure 3A**). In line with this observation, we identified high proportions of donor-specific and shared clonotypes with a fixed CDR3α sequence length of 12 amino acids, consistent with known molecular requirements for binding of the MR1-5OP-RU complex^15,41^ (**Figure 3B**). In addition to TRAV1-2+ clones, two TRAV1-2-clonotypes were identified as being shared between donors (**Figure 3C**). This included a TRAV16-TRAJ28 clonotype paired with TRBV15 and a TRAV6-TRAJ38 clonotype paired with TRBV29-1. Both TRAV1-2-clonotypes had CDR3α sequences that lacked the TRAJ-associated conserved tyrosine at position 95 which has been previously identified as critical for recognition of MR1 by TRAV1-2+ clonotypes and had variable CDR3α lengths.

**Figure 3.**
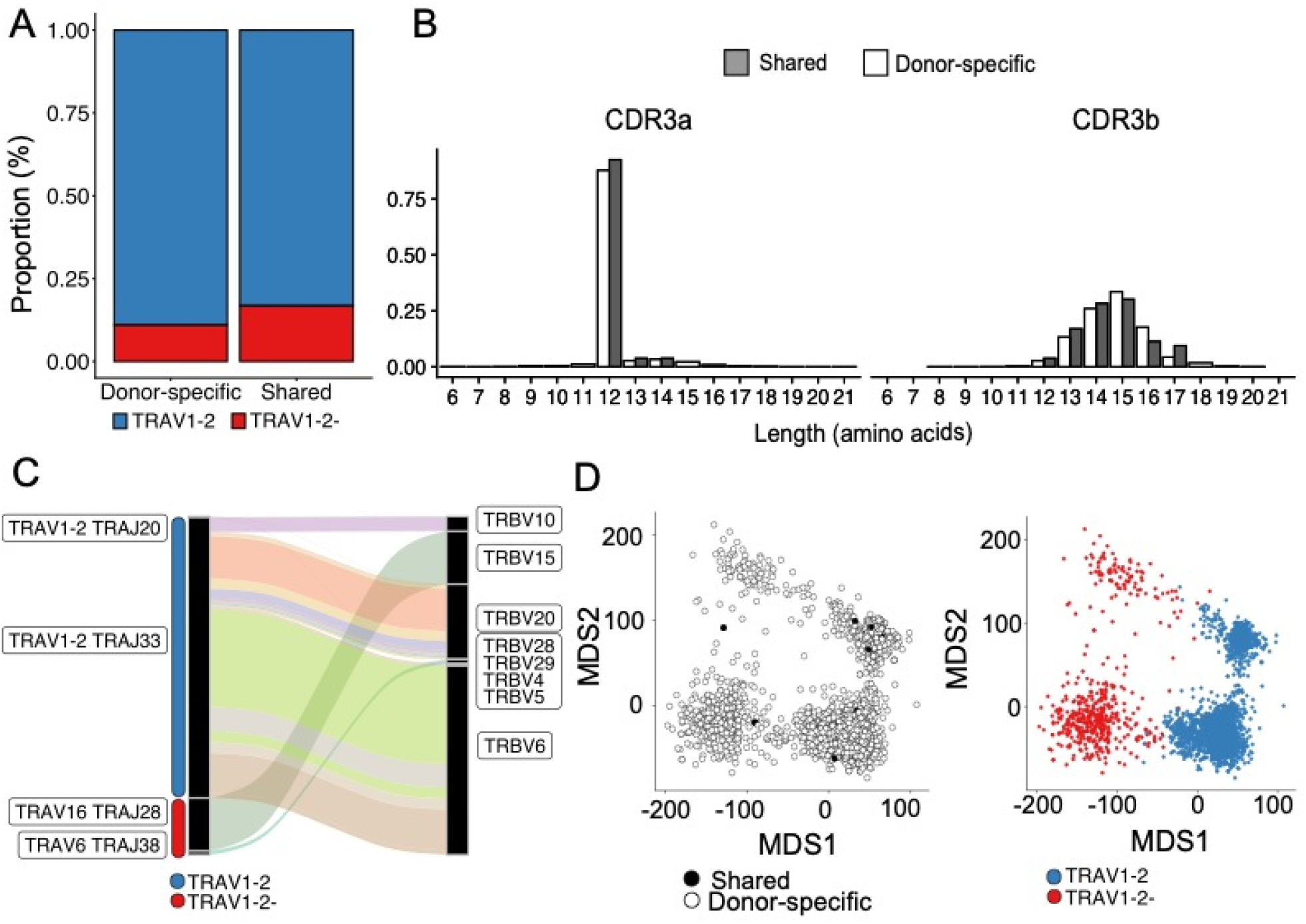
MR1T clonotype diversity is constrained by TCR gene usage and CDR3 length. (A) Bar plot showing the median proportion of TRAV1-2+ and TRAV1-2-MR1T-cells per donor (n = 8). (B) Histogram of CDR3α and CDR3β amino acid sequence lengths for shared and donor-specific clonotypes. Shaded = shared, white = donor-specific. (C) Sanky diagram showing the gene usage of shared MR1T clonotypes (n = 23 clonotypes). A clonotype was defined by nucleotide sequence and considered shared if it was identified in two or more donors. TRAV-TRAJ gene rearrangement and TRBV gene usage are labeled. The width of each connection in the plot represents the proportion of that clonotype within the shared repertoire. (D) Multidimensional scaling plot (MDS) of the TCRdist distance matrix representing the relative similarity among CDR3 sequences in 2-dimensional space (n = 5150 clonotypes). Each dot represents one sequence and is coloured by TRAV1-2 gene usage and whether it is donor-specific or shared.

To evaluate the similarity between CDR3 sequences in the donor-specific and shared repertoires, we clustered TCR clonotypes based on both CDR3α and CDR3β amino acid sequences using TCRdist^37^. Multidimensional scaling revealed significant differences in CDR3 similarity that were influenced by TRAV1-2 expression (**Figure 3D**). We noted that shared and donor-specific clonotypes did not cluster independently in this analysis, suggesting a high degree of sequence concordance between shared and donor-specific compartments (**Figure 3D**). Taken together, these data identify a broad array of non-TRAV1-2 MR1-restricted TCRs that are reactive with 5-OP-RU and are typically donor-restricted. Furthermore, we identify TRAV1-2+ MAIT-like TCRs that are similarly donor-restricted, although appear to have high CDR3 sequence homology with canonical MAIT TCRs.

### MR1T cells are characterized by transcriptional homogeneity at rest

To gain further insight into the phenotypic diversity of MR1T-cells, we simultaneously analyzed single-cell protein and gene expression data on sorted MR1 T cells^30^ (**Figure 4A**). After quality control filtering, a total of 18,251 cells with paired TCR-α and TCR-β chains were analyzed with an average of 1,876 cells per donor, which did not differ between groups (**Supplementary Figure 2D**, p = 0.89). Weighted nearest neighbors (WNN) analysis of protein and gene expression data revealed ten clusters^30^ (**Figure 4A**) (**Supplementary Table 2)**. Surface protein expression contributed significantly to clustering with expression of CD4 in particular driving stratification (**Figure 4B**). All donors contributed to all clusters in our analysis (data not shown). Further, WNN-defined clusters did not appear to be associated with RSTR status (**Figure 4B**). Genes associated with known MR1 T-cell functions including IFNγ, cytotoxicity, and activation had similar expression across all clusters suggesting limited functional delineation between clusters (**Figure 4D**). In keeping with this observation, transcription factors known to define MR1T-cell functional subsets in mice, RORγ (*RORC*) and T-bet (*TBX21*), were not significantly differentially expressed between clusters in our data^42,43^

**Figure 4.**
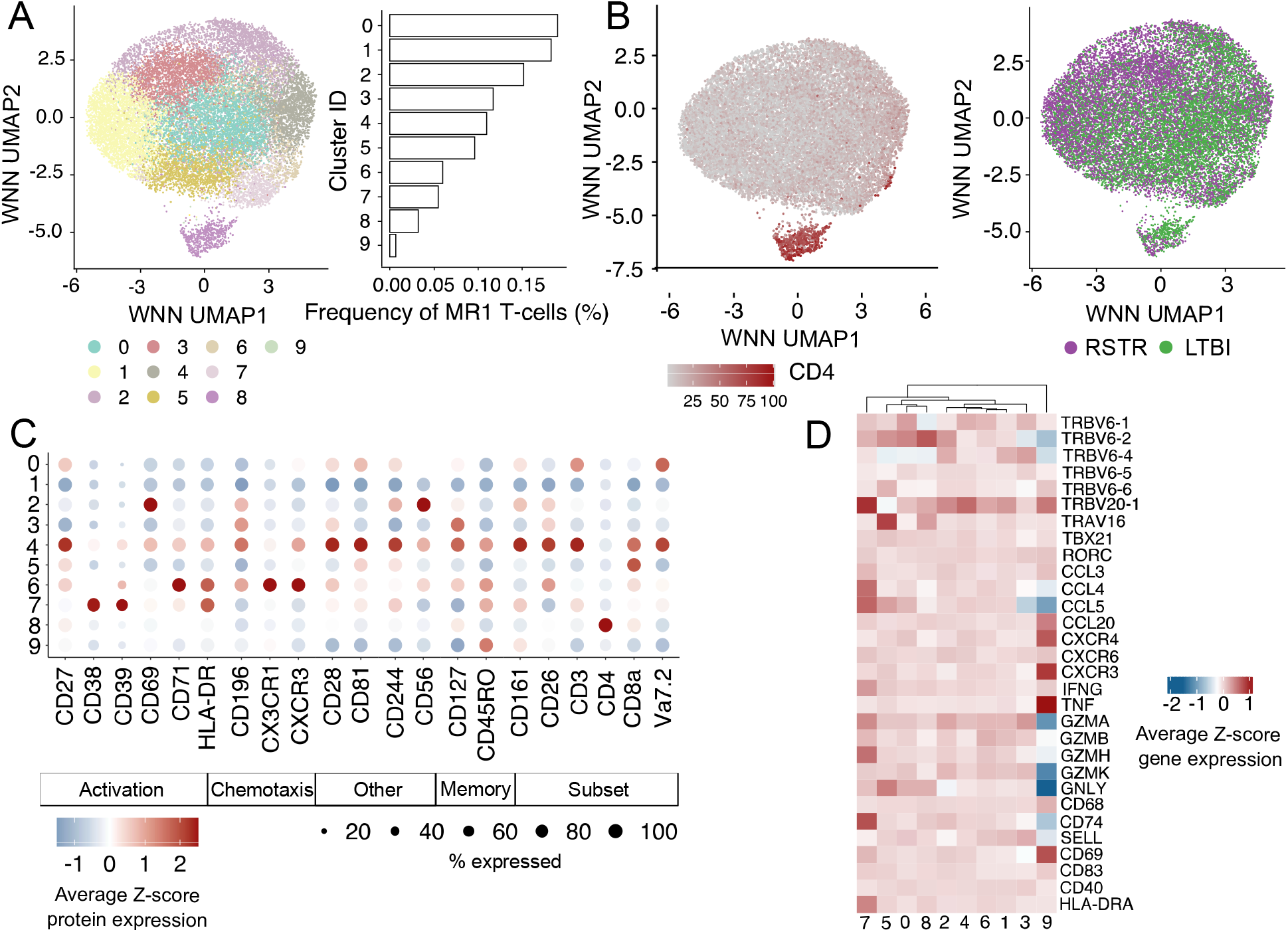
MR1T cells are characterized by transcriptional homogeneity at rest. (A) Weighted nearest neighbors (WNN) analysis was used to cluster transcriptional profiles and cell surface protein expression of 18,251 sorted MR1-5-OP-RU^+^ MR1T cells that could be annotated with both a TCR-α and TCR-β sequence. Uniform Manifold Approximation and Projection (UMAP) was used to visualize the results. The number of cell barcodes constituting each cluster is displayed in the bar chart. (B) UMAP representation of MR1T-cellss coloured by either relative expression of CD4 surface protein or TB group. (C) Dot plot showing the scaled, normalized expression of cell surface protein expression across all WNN-identified clusters. Dot size reflects the proportion of cells in a given cluster that express a marker. Dots are colored by average Z-score protein expression. (D) Heatmap of normalised, transformed gene expression of selected genes known to be associated with MR1T-cell function.

(**Figure 4D**). Surface protein expression revealed subtle differences in the magnitudes of expression of activation, memory, but the proportion of cells in each cluster expressing a particular protein was broadly similar (**Figure 4C**). Taken together, these data reveal that MR1 T cells exhibit remarkably homogeneous transcriptional programs at rest despite phenotypic and clonotypic diversity. Analysis of covariation between gene expression and TCR sequence using the clonotype neighbor graph analysis (CoNGA) package highlighted one small cluster of CD4-expressing T cells with diverse TCR sequences as having significant gene-expression/TCR correlation (**Supplementary Figure 3**). No other significant clusters were identified, consistent with overall transcriptional homogeneity.

### Immunosequencing reveals expansion of MR1T and DURT clonotypes among RSTRs

Having identified a trend towards greater MR1 clonotype sharing in RSTRs using scTCR-seq, we then sought to validate this observation in a larger set of participants in our cohort. We performed sequencing of the TCR-α and TCR-δ chains present in the peripheral blood of RSTRs (n = 19) and LTBI controls (n = 20) (**Figure 1A**). We identified a median of 80,227 and 74,915 templates per donor in RSTRs and LTBIs, respectively (p = 0.69). There was a trend toward decreased clonality among RSTRs (p= 0.065) (**Figure 5A**). We identified total 302,496 clonotypes that were shared between at least two donors representing 16.4% of the total repertoire. Notably, the distribution of TCR-□ clone sharing in this cohort, mirrored what we observed in the scTCR-Seq data with 8.01% of templates being shared between 3 or more donors (**Figure 5B**)^44^. Using two-sided Fisher’s exact tests and a nominal p-value cutoff of 0.01, we identified 303 clonotypes that were significantly associated with either RSTR or LTBI status (**Figure 5C**). Of these enriched clonotypes, two clonotypes were identical to TCR-α sequences present in sorted MR1T cells and both were preferentially detected among RSTRs (**Figure 5D**). Notably, the canonical MAIT TCR-α was detected at similar high-frequency among both RSTRs and LTBIs (**Figure 5E**). None of the MR1T clonotypes from our scTCR-Seq analysis were found to be enriched among LTBI donors (data not shown).

**Figure 5.**
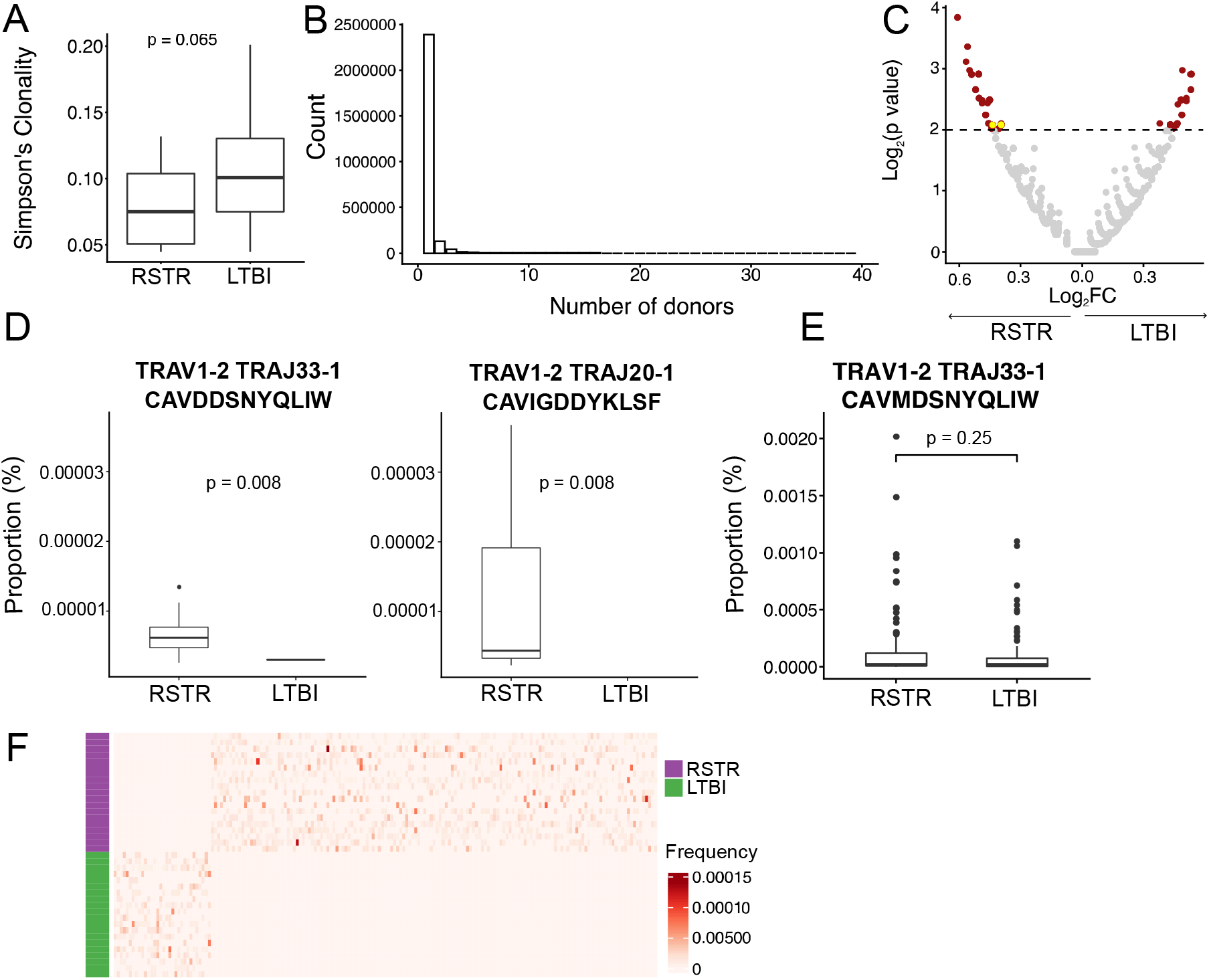
Immunosequencing reveals expansion of MR1T and DURT clonotypes that are significantly enriched among RSTRs. (A) PBMC from RSTR (n=19) and LTBI (n=20) were analyzed using TCRAD ImmunoSEQ (Adaptive Biotechnologies). Simpson’s clonality was calculated for each donor’s repertoire and stratified by group. (B) Bar plot showing the number of TCR-α/δ clonotypes that were shared between donors (n = 39) (C) Volcano plot of TCR-α/δ clonotypes after testing for enrichment in RSTR or LTBI groups. Enriched clones are highlighted in red. MR1T clonotypes identified previously in scTCR-seq are highlighted in yellow. In this analysis, a clonotype was defined by CDR3α amino acid sequence, V- and J-gene identity. Enriched clones were identified by two-sided Fisher’s exact test using a p-value threshold of 0.01 (threshold indicated by dashed line on the plot). P-values have not been adjusted for multiple testing. (D) Boxplot of frequencies of MR1T clonotypes previously defined in scTCR-seq analysis that were also identified among enriched clones in the volcano plot. Matching was based on CDR3α amino acid sequence, V- and J-gene identity. Displayed p-values were calculated using Fisher’s exact tests and are unadjusted for multiple testing (E) Boxplot of frequency of canonical MAIT TCR-α stratified by group. Displayed p-value was calculated using Wilcox Rank Sum test (F) Heatmap of frequencies (as a proportion of total templates per sample) of TCR-α/δ clonotypes found exclusively among either RSTR or LTBI donors and p < 0.05 by Fisher’s exact test. Unadjusted p-values were used to select clonotypes for inclusion in the heatmap.

Finally, we expanded our analysis to consider all possible TCR-α or TCR-δ clonotypes, irrespective of their link to MR1 T cells. Of the 303 enriched clones, we found that 147 and 33 clones were exclusively detected among RSTRs or LTBI controls, respectively (**Figure 5F**). We noted that significantly more clonotypes were nominally associated with RSTRs than would be expected by chance alone (p < 0.0001, χ2 test). Taken together, these data reveal associations between defined MR1T clonotypes as well as other clonotypes of unknown specificity and ‘resistance’ to M.tb infection.

## DISCUSSION

‘Resistance’ to M.tb infection may be mediated through both innate and adaptive immune mechanisms that are not mutually exclusive^45^. Since DURTs have features of both innate and adaptive immunity, we hypothesized that they might be associated with this unique clinical phenotype^46^. Our finding that peripheral blood frequencies of MR1T cells were higher among RSTRs led us to analyze this DURT subset in much more detail than previously possible using multi-modal scRNA-Seq. We found that most MR1T clonotypes were restricted to a single donor but clone sharing overall appeared to be more common among RSTRs. An expanded analysis of the TCR-α repertoire revealed two MR1 clonotypes as well as several other DURT clonotypes of unknown specificity that were also associated with ‘resistance’ to M.tb infection. Taken together, our results highlight the association between total MR1T cell frequency as well as specific MR1T clonotypes with ‘resistance’ to M.tb infection.

The prevailing paradigm that MR1-restricted T cells are donor-unrestricted derives from extensive studies of MAIT cells, which express a semi-invariant TCR^9^. However, the universe of MR1-restricted T cells likely extend beyond MAIT cells. A recent study used CRISPR–Cas9 screening to establish MR1-mediated killing of most human cancer types by a cytolytic T cell clone that did not express a canonical MAIT TCR. Further, concentration-dependent addition of vitamin B-related ligands of MR1 reduced TCR-mediated recognition of cancer cells, suggesting that there may be cancer-specific metabolites that are surveyed by MR1^47^. In a similar vein, MR1-restricted T cell clones expressing diverse TCRs have been shown to differentially recognize bacterial and fungal pathogens^21,22,39^. Pathogen-specific expansion of MR1-restricted clonotypes has been reported in the context of acute *Salmonella* Paratyphi A infection^40^. Mycobacterial photolumazines are MR1-presented ligands that are more potent stimulators of some MR1T cell clones than 5-OP-RU^24^. In this context, the immense clonal diversity that we observed among sorted MR1T cells may not be so surprising. At least one other study comparing TCR repertoires of MAIT cells in the blood and liver of four healthy donors using scTCR-Seq noted a similar pattern of diversity that was largely donor-restricted^48^. Though their transcriptional profiles appear to relatively homogeneous, diverse TCRs may allow MR1Ts to control the time and place they are activated in a manner that is analogous to MHC-restricted T cells.

Data from animal models have revealed conflicting evidence of a protective role of MR1 T-cells in vivo. MR1^-/-^ mice have comparable survival with WT mice following *M.tb* infection^49^. Vaccination of mice with 5-OP-RU failed to protect against subsequent M.tb challenge despite robust activation and expansion of MAIT cells in the lung^49–51^. Intratracheal administration of 5-OP-RU in non-human primates led to functional MAIT cell exhaustion rather than expansion^52^. However, 5-OP-RU treatment of mice during the chronic phase of infection was effective in reducing the bacterial burden in an IL-17A dependent manner^49^. Intravenous BCG vaccination of non-human primates was associated with sterilizing protection against M.tb challenge and robust expansion of MAIT cells in the lungs when compared to aerosol or intradermal BCG vaccination^53,54^. None of these studies sought to evaluate the association of MR1T cells and protection from M.tb infection rather than disease. In light of the clonotypic diversity that we have observed among MR1T cells, one interpretation of these data is that targeting MAIT cells that recognize 5-OP-RU may not be desirable. Rather, a diversity of MR1T cells recognizing one or more mycobacterial ligands may be recruited in the setting of BCG vaccination and mediate a superior protective effect, either directly or indirectly.

Our results must also be considered in the context of other household contact studies of MR1T cells. A cross-sectional study from Peru found that peripheral blood MAIT cell frequencies were comparable between IGRA+ and IGRA-household contacts of active TB cases^55^. A study from the Gambia found that household contacts that failed to convert IGRA over six months had lower levels of IL-17+ and IFNγ+ CD8 MAITs cells when compared to IGRA converters^56^. Finally, a cross-sectional study from Haiti revealed increased CD25 expression and Granzyme B production after in-vitro stimulation of MAIT cells from IGRA-household contacts when compared matched IGRA+ controls^57^. Despite the significant heterogeneity in study design, timing of sampling, and definition of MR1T cells, these data suggest associations between MAIT cell functional profiles and ‘resistance’ to M.tb infection that are not inconsistent with what we have observed. In contrast to the published studies, we sought to define functional profiles of MR1T cells directly ex vivo using multi-modal scRNA-Seq in the absence of in-vitro stimulation. Our data revealed known functions of MAIT cells, including expression of tissue-homing and cytotoxic markers^9^. However, we also discovered an unexpected diversity of MR1T clonotypes and associations with RSTRs that were not addressed by prior studies.

In summary, our findings advance fundamental knowledge of MR1T cell clonotypic diversity in the context of a unique longitudinal human cohort. The specific associations we report with ‘resistance’ to M.tb infection in humans are notable in light of several negative studies of MAIT cells in animal models of M.tb disease. Future work might focus on developing animal models of tuberculosis ‘resistance’ in which the role of MR1T cells can be probed more specifically^58^. At the same time, significant effort will be required to discover new MR1 ligands and determine whether antigen-specific MR1T responses can mediate clearance of M.tb and underlie the RSTR phenotype^24^.

## ACKNOWLEDGEMENTS

This work was supported by the U.S. National Institutes of Health (R01-AI124348 to Boom, Stein, and Hawn; R01-AI125189 and R01-AI146072 to Seshadri) and the Bill & Melinda Gates Foundation (OPP1151836 and OPP1109001 to Hawn and GH-VAP-IS-ID5 to Seshadri). We thank Dr. Branch Moody and Professor Adriaan Minnaard for providing lipid antigens. Additionally, the authors would like to acknowledge The Fred Hutch Genomics Core for sequencing and Rachel Gittelman and Emma Bishop for assistance with the computational analyses. Finally, we acknowledge the NIH Tetramer Core Facility for providing tetramers and monomers used in this study. The Tetramer Core Facility is supported by contract 75N93020D00005 from the National Institute of Allergy and Infectious Diseases, a component of the National Institutes of Health in the Department of Health and Human Services.

We also appreciate the efforts of the Uganda-CWRU Research Collaboration team that was involved in recruitment, clinical characterization, and data management for the cohort: Drs. Mary Nsereko and Ronald Kiyemba, LaShaunda Malone, Keith Chervenak, Sophie Nalukwago, and Hussein Kisingo.

## AUTHOR CONTRIBUTIONS

D.L.C. performed the data integration, analysis, and visualization. D.L.C. and C.S. wrote the paper with contributions from all authors. Flow cytometry data were acquired by E.D.L. and K.K.QY. and M.S.A. performed MR1T-cell sorting. K.K.Q.Y, M.S.A. and S.L. performed multi-modal single-cell RNA sequencing. H.M.K, C.M.S, T.R.H. and W.H.B facilitated access to clinical specimens. E.N. oversaw scRNA-Seq and advised on MR1T sorting strategy. P.B performed CoNGA analysis including visualization.

## FIGURES

**Supplementary Figure 1.**
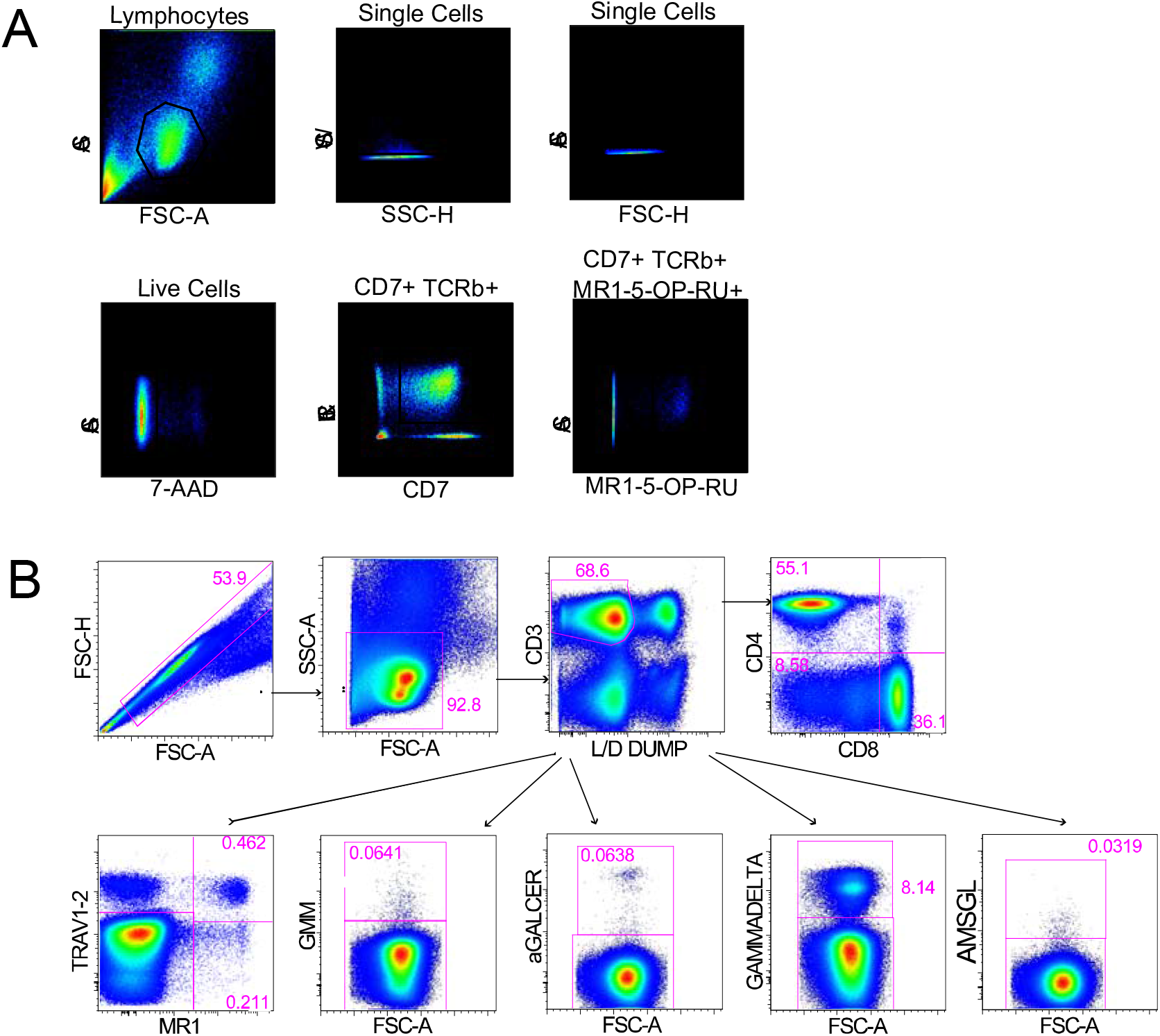
(A) Gating scheme identifying donor-unrestricted T-cells in peripheral blood of RSTR and LTBI donors (B) Sorting scheme identifying CD7+ TCRab+ MR1-5-OP-RU+ cells in peripheral blood of RSTR and LTBI donors.

**Supplementary Figure 2.**
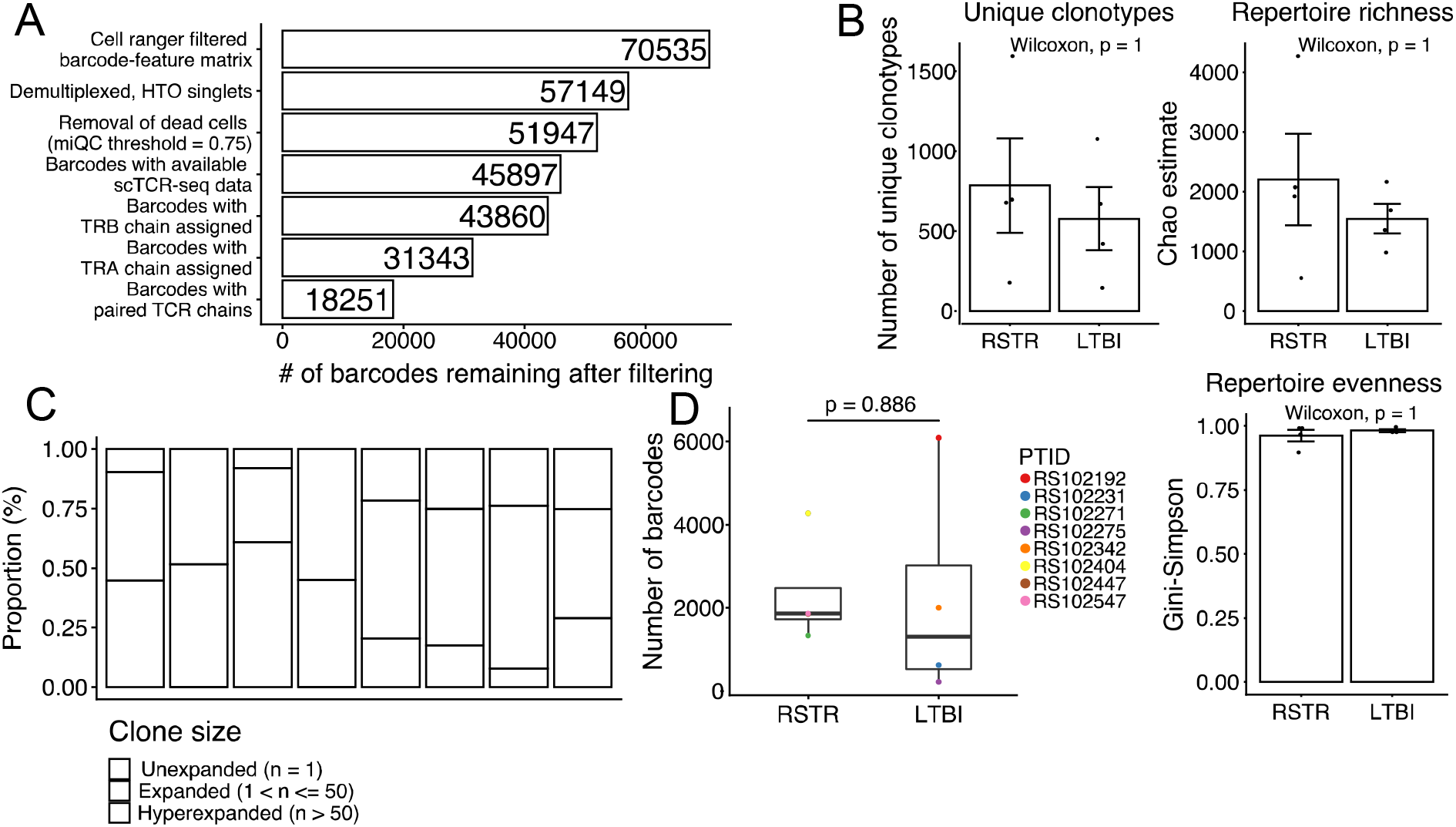
(A) Summary of CITE-seq QC filtering (B) Global diversity metrics summarizing the MR1T repertoire. (C) The proportion of unexpanded (n = 1), expanded (1<n<=50) and hyperexpanded (n > 50) MR1T clonotypes within each donor. A clonotype is defined based on nucleotide sequence (D) The number of barcodes per sample, compared between TB groups. Statistical testing was performed using a Wilcoxon Rank Sum test.

**Supplementary Figure 3.**
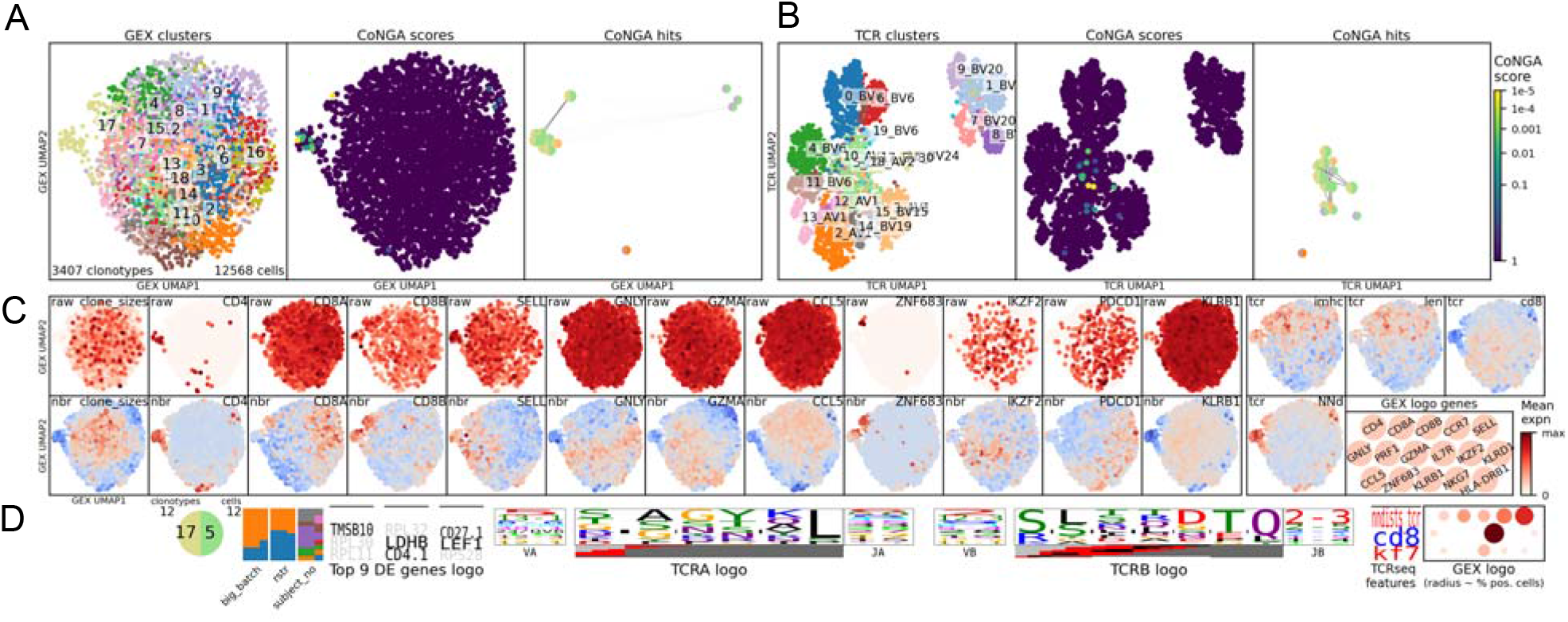
Graph-vs-graph analysis of MR1T clonotypes (A) UMAP of MR1 T clonotypes based on similarity in gene expression. Panels are coloured left to right by gene expression cluster, CoNGA score – indicating the clonotypes with significant overlap between their gene expression and TCR similarity – and CoNGA hits (clonotypes with a significant CoNGA score). (B) UMAP of MR1 T clonotypes based on similarity in TCR CDR3 sequence. Panels are coloured left to right by TCR cluster, CoNGA score and significant CoNGA scores. (C) gene expression UMAP colored by gene expression (D) Consensus logos for CDR3α and CDR3β sequences, V and J gene usage for all clones with a significant CoNGA score.

